# Predictive Gene Discovery with EPCY: A Density-Based Alternative to DE analysis

**DOI:** 10.1101/2025.08.07.668357

**Authors:** Eric Audemard, Jean-François Spinella, Vincent-Philippe Lavallée, Josée Hébert, Guy Sauvageau, Sébastien Lemieux

**Affiliations:** The Leucegene project at Institute for Research in Immunology and Cancer, Université de Montréal, Montréal, Québec, H3T 1J4, Canada; Centre Hospitalier Universitaire Sainte-Justine Research Center, Montréal, Québec, H3T 1C5, Canada; Quebec Leukemia Cell Bank, Maisonneuve-Rosemont Hospital, Montréal, Québec, H1T 2M4, Canada; Department of Biochemistry and Molecular Medicine, Faculty of Medicine, Université de Montréal, Montréal, Québec, H3T 1J4, Canada; Department of Medicine, Faculty of Medicine, Université de Montréal, Montréal, Québec, H3T 1J4, Canada; Division of Hematology-Oncology, Maisonneuve-Rosemont Hospital, Montréal, Québec, H1T 2M4, Canada; Department of Pediatrics, Faculty of Medicine, Université de Montréal, Montréal, Québec, H3T 1J4, Canada; Hematology and Oncology Division, Centre Hospitalier Universitaire Sainte-Justine, Montréal, Québec, H3T 1C5, Canada; Mila – Quebec AI Institute, Montréal, Québec, H2S 3H1, Canada

## Abstract

Identifying predictive genes from high-throughput data remains a key challenge in biomedical research. Most current approaches rely on statistical tests to select differentially expressed genes (DEGs), which may not align with the goal of predicting outcomes. We present EPCY, a method that ranks genes based on their predictive power using cross-validated classifiers and density estimation, without relying on null hypothesis testing. Applied to both bulk and single-cell RNA sequencing datasets, EPCY consistently outperforms benchmark DEG-based methods in selecting robust candidate genes. It also demonstrates greater stability across varying cohort sizes, enabling reproducible gene prioritization even in large, heterogeneous datasets. EPCY provides interpretable predictive scores, facilitating candidate selection aligned with downstream validation goals.

## Introduction

In medical research, associating a predictive signal with a clinical outcome—i.e., identifying a gene expression to use as a biomarker—is a critical step toward elucidating disease mechanisms, improving patient stratification, and ultimately optimizing treatment strategies. Large-scale transcriptomic datasets from complex diseases such as cancer (1), in combination with statistical methods for differential expression (DE) analysis between defined sample groups (e.g., DESeq2 (2), Limma-Voom (3), EdgeR (4)), are commonly used as an initial approach to identify putative candidate genes associated with specific conditions, subgroups, or clinical outcomes.

Significant differentially expressed genes (DEGs) are typically further filtered using integrative analyses based on prior biological knowledge, such as gene function or pathway membership (1,5), to retain a smaller set of candidates for downstream validation. Although such approaches have led to the identification of some effective predictive genes (PGs) (6)—i.e., DEGs that also satisfy predictive performance criteria—the underlying statistical framework is not designed to optimize for predictiveness. As a result, these methods often yield suboptimal results, characterized by reduced sensitivity and specificity.

Furthermore, candidate gene identification is often conducted using large cohorts (>100 samples), such as those from TCGA, BEAT AML (7) or Leucegene (8) which exhibit substantial inter-sample variability—even within ostensibly homogeneous molecular subgroups. In contrast, most DE methods were developed and optimized for small-scale studies with limited intra-group variance, where highly variable genes are typically filtered or penalized to control the false discovery rate (2–4).

Here, we introduce EPCY, a general density-based method that ranks genes based on the predictive capacity of their expression profiles, without relying on null hypothesis testing. Instead, EPCY leverages cross-validation (CV), a standard machine learning approach (9), to assess performance on unseen samples.

Using both bulk and single-cell RNA-Seq data—derived from in-house Leucegene and publicly available datasets—we demonstrate that EPCY outperforms benchmark DE methods, including DESeq2 (2), EdgeR (4) and limma voom (3) for bulk sequencing data, as well as Limma Trend (3), MAST (10) and the Wilcoxon rank-sum (the default Seurat (11)) for single-cell data, in identifying high-confidence predictive gene candidates. Notably, we demonstrate that EPCY provides interpretable and reproducible results in large cohorts—including datasets with >100 samples or cells—whereas standard DE analyses often lose discriminatory power as sample size increases.

## Results

### Method Overview

EPCY evaluates the predictive capacity of each gene individually and returns corresponding predictive scores along with their confidence intervals. To ensure the reliability of these scores, EPCY employs a leave-one-out cross-validation scheme, training gene-specific Kernel Density Estimation (KDE) classifiers to assess performance on held-out samples. A schematic overview is provided in Figure 1A, and implementation details are described in the Methods section.

**Figure 1.**
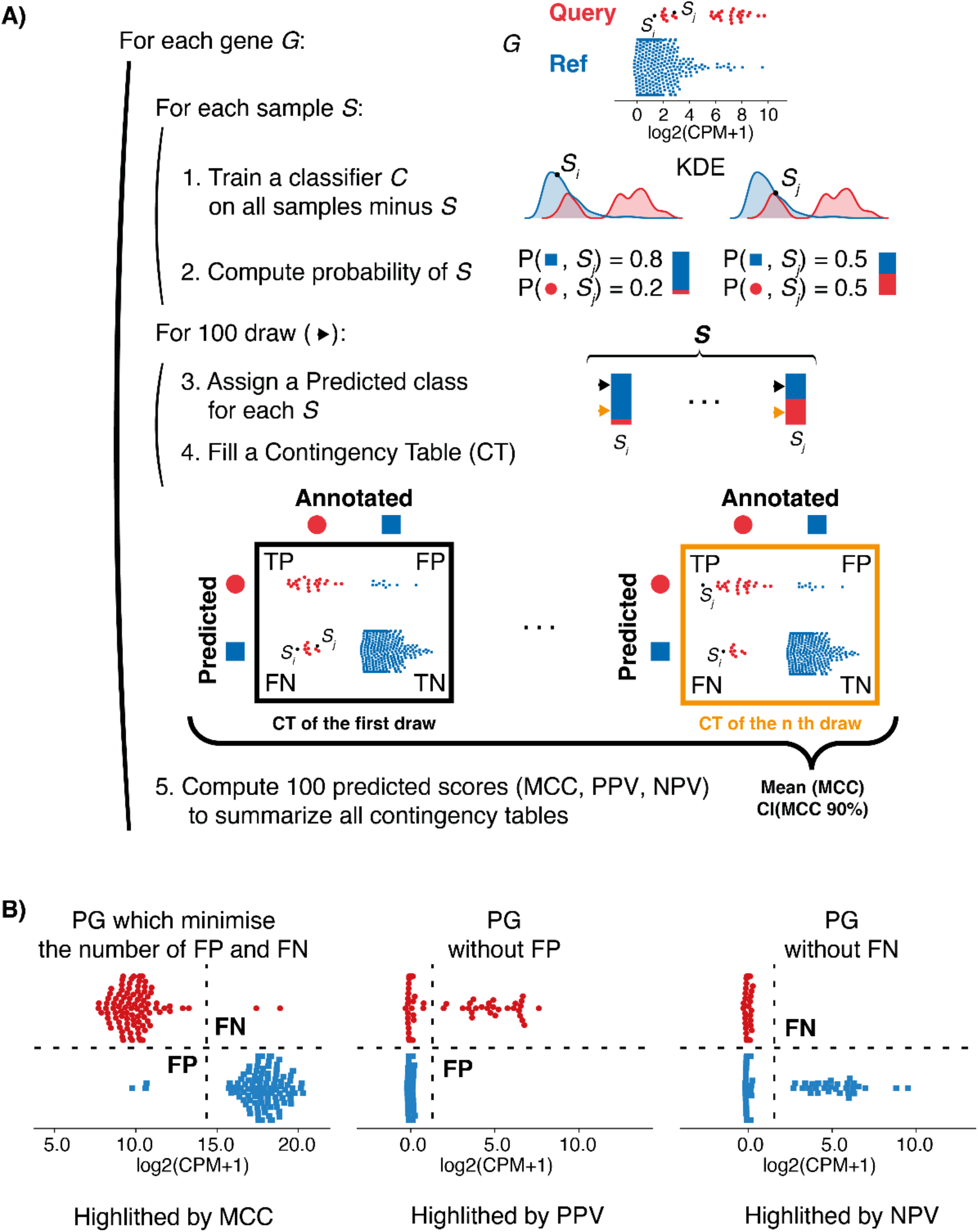
A) Schematic overview of the EPCY method. For each gene, EPCY evaluates predictive capacity using a leave-one-out cross-validation approach. Subgroup densities are modeled using Kernel Density Estimation (KDE), and a KDE-based classifier *C*_*kde*_ is trained on all samples except one. Class probabilities are then predicted for the held-out sample. This process is repeated for all samples. For each sample, a predicted class is drawn based on the classifier’s probability distributions, and a contingency table (CT) is filled. This sampling is repeated *m* times (default *m* = 100) to generate multiple CTs. Matthews Correlation Coefficient (MCC) is computed for each CT, and the final predictive score is reported as the mean MCC with a 90% confidence interval (CI). B) EPCY can also compute other predictive metrics from the CTs, including Positive Predictive Value (PPV) and Negative Predictive Value (NPV), which are particularly useful in single-cell datasets where dropout is frequent.

Although this cross-validation approach enhances performance assessment, it is more computationally intensive than conventional differential expression (DE) methods due to repeated training of KDE classifiers. The computational complexity of the algorithm is *O*(mn^2^p), where *m* is the number of draws, *n* the number of samples, and *p* the number of genes. For example, using the current implementation with four threads (Intel E5-2640 v4), analyzing the Leucegene cohort (655 samples, 60,564 annotated genes) requires approximately 1.5 hours, compared to under 10 minutes for standard DE methods (see Table S1). However, this runtime is generally acceptable for gene prioritization tasks, and various strategies could be applied to accelerate the implementation if needed.

By default, EPCY reports the Matthews Correlation Coefficient (MCC), selected for its informative evaluation of binary classification and robustness to class imbalance (12). The MCC ranges from −1 to +1, with +1 indicating perfect prediction, 0 indicating random performance, and −1 indicating complete disagreement.

Notably, both the KDE classifier and the predictive scores used to summarize contingency tables can be customized to target specific quantification profiles. This is particularly useful in single-cell datasets, where Positive Predictive Value (PPV) and Negative Predictive Value (NPV) can be employed to identify predictive genes that minimize false positives or false negatives, as illustrated in Figure 1B. These metrics are especially relevant in the context of high dropout rates. Additional predictive metrics, such as F1-score or accuracy, are available and described in the EPCY documentation.

### Impact of cohort size on genes selection

Differential expression (DE) methods were originally designed for comparisons involving small sample sizes. To evaluate how both DE algorithms and EPCY perform as cohort size increases, we conducted a systematic analysis using the relatively homogeneous t(15;17) AML subgroup from the Leucegene dataset (n = 30) compared to other AML samples (n = 625). We generated 10 different cohort designs by subsampling with replacement, each replicated 10 times to assess variability.

Our results show that as cohort size increases, DE methods tend to produce more significant p-values, which leads to highly variable gene candidate lists when fixed p-value thresholds are applied. Moreover, the standard deviation of these p-values across replicates also increases, reducing the reproducibility of DE results (Figure 2A). In contrast, EPCY rapidly stabilizes its Matthews Correlation Coefficient (MCC) score once enough samples are reached, enabling consistent candidate selection based on a fixed MCC threshold, regardless of cohort size.

**Figure 2.**
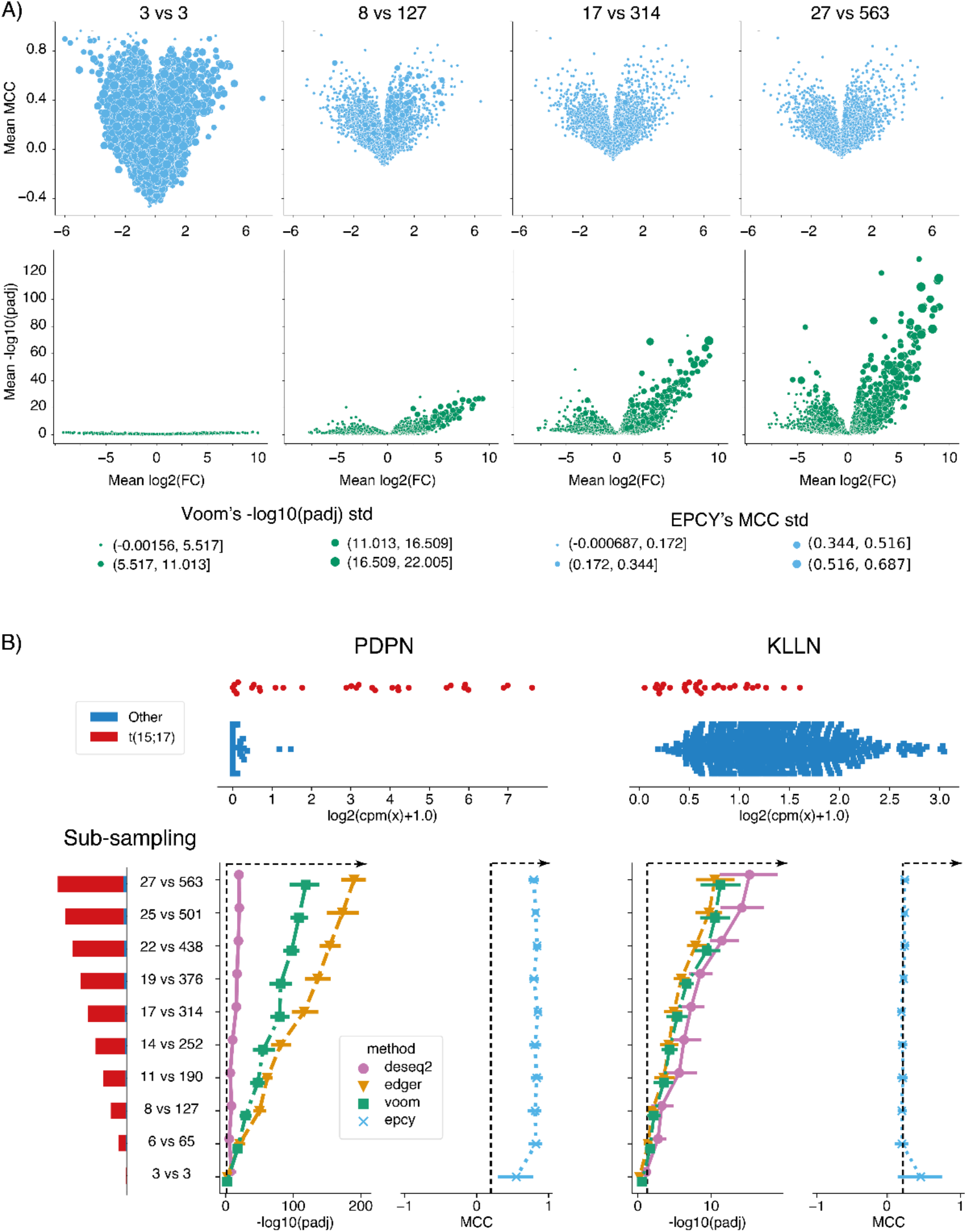
Illustrates trends of EPCY’s MCC and adjusted p-values (padj) from DE methods, as a function of the number of samples considered. For this purpose, the Leucegene cohort is randomly sub-sampled to simulate 10 replicates of 10 different cohort sizes, with comparisons ranging from 3 vs 3 to 27 vs 563 and adding 10% samples at each step. Panel A Volcano plots showing the relationship between adjusted p-values (padj) from DE methods and MCC scores from EPCY across increasing cohort sizes. DE methods show increasing padj variability and significance with larger cohorts, while EPCY’s MCC scores stabilize early and remain consistent. Panel B) use expression profiles of two genes, PDPN and KLLN, illustrate how DE methods may prioritize genes with overlapping expression across groups, while EPCY maintains robustness by focusing on predictive separation. Error bars represent 95% confidence intervals across 10 replicates per cohort size. Dashed lines indicate common selection thresholds: padj < 0.01 and MCC > 0.2.

Importantly, DE methods exhibit biases not only for genes with clear differential expressions (e.g., *PDPN* in t(15;17) AML) but also for genes with overlapping expressions across groups (e.g., *KLLN*, Figure 2B). EPCY, by contrast, maintains robustness in both scenarios.

It is worth noting that EPCY may return negative MCC values when applied to very small sample sizes (e.g., 3 vs. 3), due to insufficient data for KDE classifiers to model expression distributions accurately. These negative scores serve as a useful diagnostic for underpowered analyses. While no universal minimum sample size can be defined, since it depends on data heterogeneity, our results suggest that using at least 50 samples, with a minimum of 10 per subgroup, significantly reduces the occurrence of negative MCCs. Alternatively, tuning KDE parameters (e.g., bandwidth) can improve robustness in smaller datasets. Nevertheless, EPCY has been designed to analyze large cohorts and analysis of small cohorts should be performed with caution.

Additionally, EPCY computes log2 fold changes directly from raw expression values, whereas DE methods estimate fold changes from model-fitted distributions. This methodological difference explains the discrepancies observed between EPCY and DE fold change values (Figure 2A and Figures S1).

### Predictive Gene versus Differentially Expressed Genes

Candidate gene selection often relies on arbitrary thresholds (e.g., p-values or MCC scores), whose distributions are highly dependent on the experimental design. To fairly compare EPCY with standard differential expression (DE) methods, we selected a fixed number of top-ranked genes based on either the lowest p-values or highest MCC scores and then compared their expression profiles. Applying this approach to the t(15;17) subgroup of the Leucegene cohort revealed substantial discrepancies between EPCY and conventional DE tools (Figure 3).

**Figure 3.**
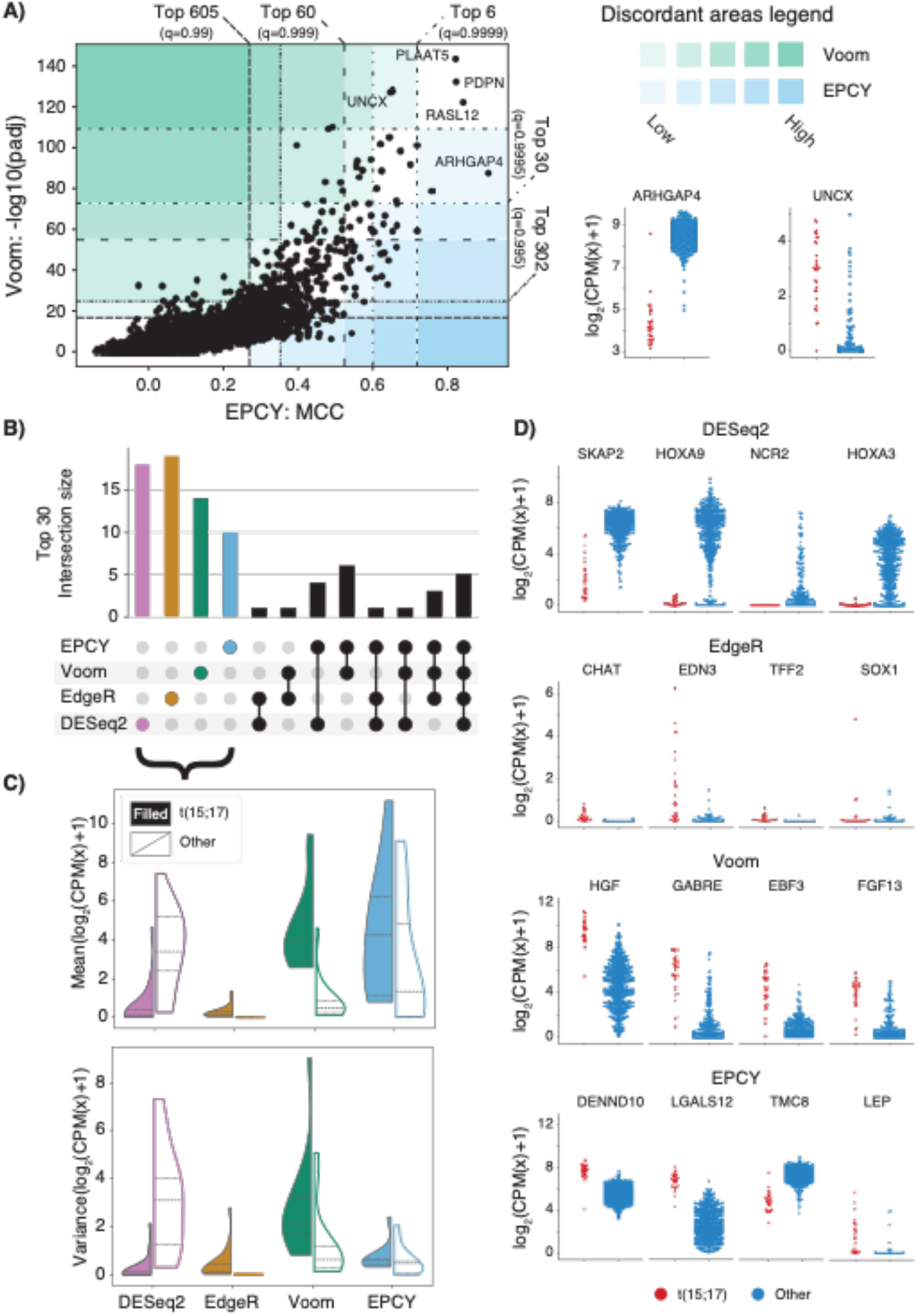
Comparison of EPCY analysis of t(15;17) AML samples versus other AML samples of the Leucegene cohort to other differential expression analysis methods. Panel A) shows adjusted p-values (padj) derived from the Limma Voom alongside EPCY’s Matthews Correlation Coefficient (MCC) for all genes. Dashed lines represent thresholds based on quantiles for each method, highlighting areas where PG and DEG are discordant. Expression profiles of ARHGAP4 (EPCY-favored) and UNCX (Limma-favored) illustrate differences in overlap between groups. Panel B) displays the number of intersecting genes within top 30 for each method. Empty intersections are omitted. Further intersections using alternative thresholds can be found in supplementary data (Figure S3, S4 and S5). Panel C) presents the density of mean expression (top panel) and variance (bottom panel) for mutually exclusive selections of genes. Quartiles are indicated by vertical dashed lines. See supplementary data for alternative thresholds. Panel D) illustrate expression profiles of the top four uniquely selected genes per method, highlighting method-specific biases.

For instance, Figure 3A highlights the contrasting prioritization of genes by EPCY (via MCC) and Limma Voom (via adjusted p-values). Genes such as *ARHGAP4* (favored by EPCY) and *UNCX* (favored by Limma Voom), illustrate EPCY’s capacity to favorize candidates with minimum overlap between samples. These discrepancies are even more pronounced with DESeq2 and EdgeR (see Figure S2), where some top-ranked genes (top 6) show very low MCC scores (< 0.2), indicating poor predictive power.

Figure 3B shows that only 5 out of the top 30 genes are shared across all DE methods, while at least 14 are unique to a single method. Figures 3C and 3D further illustrate that these unique genes display distinct expression profiles depending on the method: Limma Voom tends to select highly expressed genes with low variance, DESeq2 favors lowly expressed genes, and EdgeR highlights genes with low overall variance. These biases reflect the statistical assumptions of each method (e.g., Poisson or negative binomial models), which may penalize atypical expression profiles such as that of PDPN using DESeq2 (see Figure S2).

In contrast, EPCY directly assesses the overlap of expression profiles between groups using the MCC, offering a more balanced and less biased evaluation. It identifies 20 genes shared with at least one DE method (Figure 3B), and the genes uniquely selected by EPCY do not cluster around a single expression pattern (Figures 3C and 3D). This density-based approach, grounded in contingency tables, is more robust to outliers and intra-group heterogeneity.

This straightforward feature of PG should be an edge to select and explore candidate genes, avoiding misleading interpretations compared to DEGs adjusted p-values. Particularly when interested in highly or lowly expressed genes (or those showing a large amplitude between conditions), setting a cutoff on (absolute) fold change is a more readable way than select a method which promote these genes indirectly. Especially when analyses are presented and summarized by a volcano plot (see Figure 1A).

### Single-cell interpretation and reproducibility

Challenges encountered in differential analyses of bulk RNA-seq cohorts are likely to extend to other forms of quantitative data, particularly single-cell datasets, where compared groups often consist of hundreds or thousands of individual cells.

To demonstrate this, we used a 10X dataset composed of 10 PBMC cell types selected by FACS and sequenced separately (13). By aggregating these 10 experiments, we obtained a dataset of

∼95,000 cells with ∼25,000 reads per cell. Unlike recent single-cell datasets, this one benefits from a cell labeling method independent of gene expression profiles, which was used as input for both differential and EPCY analyses. To assess the reproducibility of each method in distinguishing each cell type from all others, we generated 20 replicates datasets that mimic typical single-cell experiments. Each replicate consisted of ∼10,000 cells, sampled from a normal distribution centered at 10,000 with a standard deviation of 1,000. Additionally, to assess the influence of cohort size on reproducibility, we repeated the experimental setup using datasets with varying mean cell counts—3,000, 5,000, and 8,000 cells.

As observed in bulk sequencing (Figure 2A), increasing the number of cells leads to greater variability in p-values computed by benchmark algorithms like MAST. In contrast, EPCY leverages the increased sample size to stabilize the predictive score for each gene (Figure 4A). Notably, the reduced variability of genes with high -log10(padj) (> 280) is largely due to technical limitations that set adjusted p-values < 1e-324 to 0. Although the standard deviations (SD) of MCC values from single-cell and bulk RNA-seq data are not directly comparable, it is reassuring that the highest SD observed in single-cell data remained lower than in bulk RNA-seq. This is expected given the larger sample size and suggests that EPCY is robust to the noise introduced by higher dropout rates due to low sequencing coverage.

**Figure 4.**
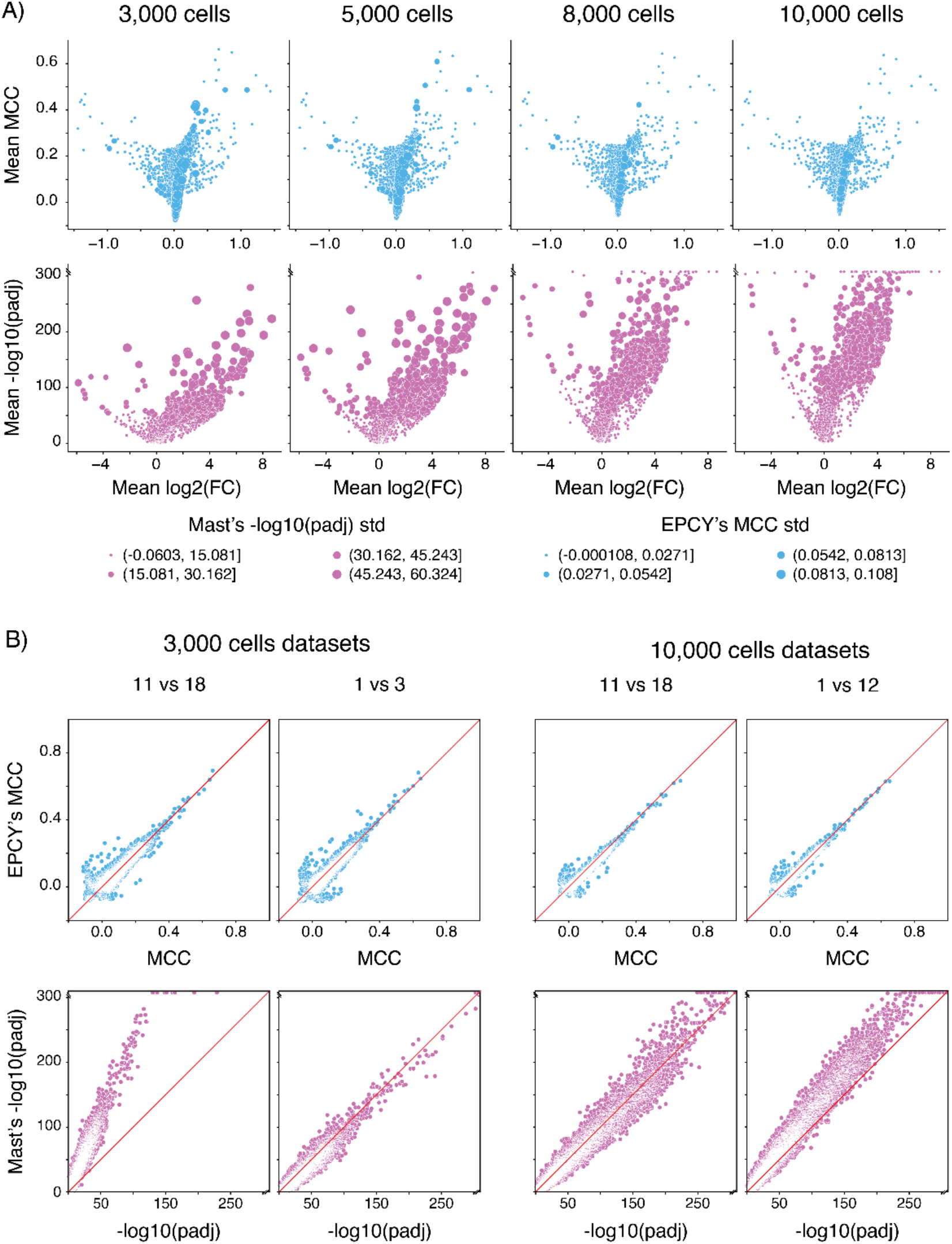
Comparison in the reproducibility of MAST and EPCY in distinguishing CD34-possitive cells from other peripheral blood mononuclear cell (PBMC) types, using 20 replicates of different sizes (3,000, 5,000, 8,000 or 10,000 cells) generated by of random sub-sampling of single-cell RNA datasets derived from a large 10X dataset. Comparison with Limma Trend and Wilcox adjusted p-values are in supplementary. Panel A) Volcano plots illustrate the relationship between MAST adjusted p-values (padj) and their standard deviations across the 20 replicates, highlighting how both metrics increase with cohort size. In contrast, EPCY’s Matthews Correlation Coefficient (MCC) remains largely stable, showing minimal variation regardless of the number of cells. Panel B) Scatter plots of adjusted p-values and MCCs resulting from the analysis of two pairs of replicates constituted of 3,000 and 10,000 cells, respectively. The red diagonal line indicates the expected trend for perfect reproducibility between datasets. Of note, using MAST 3,000 cells is sufficient to reach the technical limitation which set p-values to 0 in the 11^th^ replicate.

As observed in bulk sequencing (Figure 1A), increasing the sample size (here, the number of cells) results in greater variability of p-values when calculated using benchmark algorithms like MAST. In contrast, EPCY leverages the increased sample size to stabilize the predictive score for each gene (Figure 3A). Of note, the reduced variability of genes with high -log10(padj) (> 280), is largely due to the technical limitation that set adjusted p-values < 1e-324 to 0. Additionally, although the standard deviations (SD) of MCC values computed from single-cell and bulk RNA-seq data are not directly comparable, it is reassuring to note that the highest SD observe on single cell remained lower than that in bulk RNA-seq. This is expected given the larger sample size in single-cell experiments and suggests that EPCY is resistant to the noise introduce by higher dropout rates, linked to the low sequencing coverage.

Interestingly, while EPCY showed stable MCC scores across simulated replicates (Figure 4B and Figure 5A), we observed significant variability in p-values using MAST (Figure 4B) and other methods (Figures S6 and S7). This variability is reflected in the slope of the scatter plots and the Pearson correlation coefficient, which are harder to mitigate. As shown in Figure 5A, this issue is not limited to a specific analysis but is observed more broadly. Importantly, in all simulations, EPCY demonstrated that increasing the number of cells allows for lowering the selection threshold without compromising reproducibility. This contrasts sharply with DE methods, where the most reproducible adjusted p-values often correspond to less informative genes, consistent with their goal of eliminating non-significant profiles rather than identifying the most predictive candidates.

**Figure 5.**
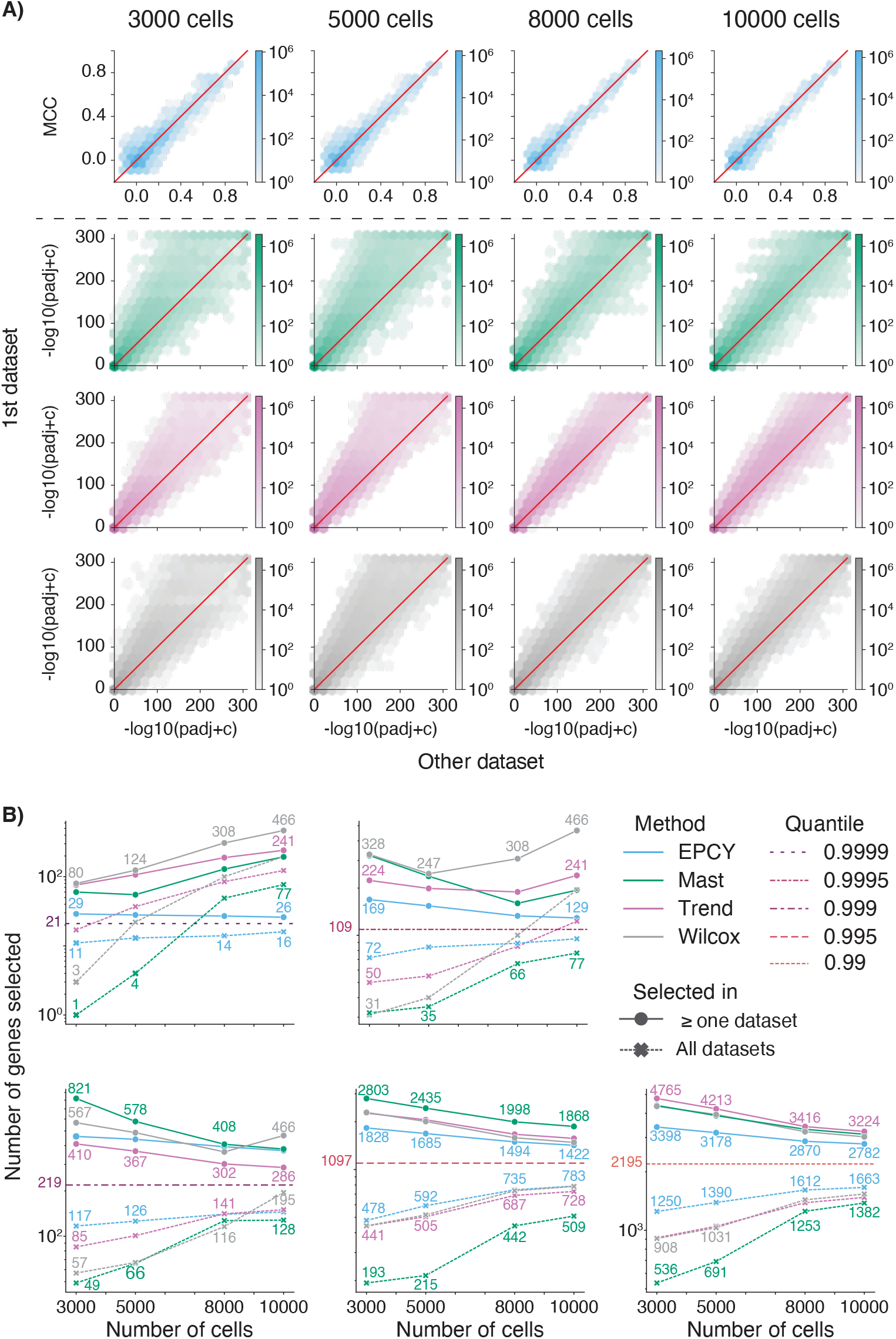
Comparison of the reproducibility of EPCY, Limma Trend, MAST, Wilcoxon analysis in distinguishing different cell types within 10 PBMC populations. Analysis is based on 20 randomly sub-sampled single-cell RNA datasets of variable sizes (3,000, 5,000, 8,000, and 10,000 cells) derived from a large 10X dataset. Panel A) compare the reproducibility of adjusted p-values and Matthews Correlation Coefficient (MCC) values computed on 20 replicates datasets. Cumulative number of genes within each hexagon area is aggregated across all pairwise comparisons between the 1^st^ dataset (y-axis) and 19 other datasets (x-axis), for all analyses aimed at distinguishing one PBMC cell type from the other nine. Red diagonal lines indicate the expected trend of a perfect reproducibility across all comparisons. Panel B) compare the reproducibility of selected candidate genes across the 20 replicates datasets analyzed, using the number of distinct candidate genes selected to discriminate each cell type in at least one (solid lines) and all replicates (dashed lines), across different dataset size. Horizontal lines represent the expected number of distinct candidate genes, depending on the quantiles, if the analyses were perfectly reproducible across all replicates.

Consequently, EPCY analyses allow researchers to assess selected genes in the context of their discovery *i*.*e*., their predictive score and the cohort size. For example, if a candidate gene has an MCC of 0.4 using 3,000 cells, it would be prudent to validate its predictive score in other datasets or increase the number of cells to 10,000 before further investigation. Conversely, a gene with an MCC ≥ 0.8 may not require additional validation (Figure 5A).

Overall, EPCY consistently outperformed other tested methods in reproducibility, even if we focus on common selected candidate genes (Figure 4B), particularly in higher quantiles where DE methods are limited by the high number adjusted p-values set to 0. Conversely, if the goal is to select a large number of genes (lower quantiles), for downstream analysis, despite a reduction in the difference between EPCY and other methods, our approach still demonstrated superior performances. This should lead to greater reproducibility in subsequent analyses, such as pathways enrichment studies.

## Discussion

Analysing large cohorts, our study highlights the limitations of DE methods in selecting a manageable and informative subset of genes for further investigation, and introduce EPCY and its Predictive Genes (PGs) as a more effective and reproductible alternative.

Recent work by Li *et al*. (14) showed that parametric statistical tests, particularly EdgeR (4) and DESeq2 (2), tend to underestimate FDR in large datasets. While non-parametric alternatives like the Wilcoxon rank-sum test have been recommended for cohorts with more than eight samples per condition. We explored this approach with the Leucegene cohort, yielding promising preliminary results. However, we argue that the low validation rate of DEGs is not solely due to FDR underestimation. Instead, it seems largely drive by an overpowered nature of statistical tests in large cohorts (like TCGA, BEAT AML, Leucegene), combined with the overinterpretation of p-values as a filtering mechanism.

As Liu *et al*. (15) noted, increasing the number of replicates improves detection of DEGs. However, this also increases the likelihood of detecting statistically significant but biologically irrelevant. In line with Venet *et al*. (16), who showed that 90% of random signature > 100 can predict outcomes, we suggest that gene expression differences are nearly always detectable with enough samples, making null hypothesis rarely valid. It’s akin to increasing screen size without enhancing resolution or zooming out to provide a wider shot without adjusting focus.

DE methods do not leverage additional samples to improve gene ranking. Instead, they force researchers to apply more stringent on p-values threshold, while fold remain largely unchanged. This reflects a disconnect between statistical rigor and biological relevance — a point echoed by John Tuckey’s famous advice: “Far better an approximate answer to the right question which is often vague, that an exact answer to the wrong question, which can always be made precise” (17).

To address the right question — identifying the most predictive genes — we introduce EPCY. It evaluates the predictive capacity of each gene expression profile and benefits from larger sample sies to refine its results. EPCY’s predictive scores are directly interpretable, based on true/false positives and negatives, and increasing stringency leads to more distinct expression profiles between groups. This contrasts with DE methods, where stringency often highlights genes that best fit statistical assumptions rather than biological relevance.

Moreover, EPCY’s scores are consistent across datasets: two genes with the same MCC will have similar predictive power, regardless of the dataset used. This consistency could help standardize gene prioritization pipelines and reduce variability across studies.

However, EPCY’s strength in large datasets comes with a caveat: caution is needed when analyzing small cohorts. In such cases, a trade-off between sequencing depth and sample size must be considered (15), depending on the expected predictive power. As shown with the 10X single-cell dataset, 3,000 cells are sufficient to identify PGs with MCC > 0.8, but genes with MCC ∼0.4 may require validation in larger datasets.

Although MCC is the default score in this study, EPCY supports other metrics (e.g., PPV, NPV), which may be more appropriate in specific contexts — such as managing dropout in single-cell data. Further exploration of these alternatives is warranted.

Finally, EPCY’s flexibility makes it applicable beyond transcriptomics. Any high-dimensional dataset using statistical tests to identify candidate markers — such as mass spectrometry data (e.g., Leucegene surfaceome (18)) — could benefit from EPCY’s predictive framework.

## Materials and methods

### Bulk RNA dataset and gene expression

This study is part of the Leucegene project, an initiative approved by the Research Ethics Boards of Université de Montréal and Maisonneuve-Rosemont Hospital. All AML samples were collected with informed consent between 2001 and 2018 according to Québec Leukemia Cell Bank procedures. RNA-Seq data were deposited in the Gene Expression Omnibus (GSE232130).

Workflow for sequencing has been described previously (19). Briefly, libraries were prepared with TruSeq RNA Sample Preparation kits (Illumina) and sequencing was performed using an Illumina HiSeq 2000 with 200-cycle paired-end reads. GSE232130 have a total of 691 samples, we remove samples 08H092 and 19H045 (suspected to have a mixed tumor) and all relapse samples resulting in a 655 primary AML sample (see Table S2 for sample IDs and annotations).

Sequences were trimmed for sequencing adapters and low quality 3’ bases using Trimmomatic version 0.38 (20) and aligned to the reference human genome version GRCh38 (gene annotation from Gencode version 32, based on Ensembl 98) using STAR version 2.7.1a (21). Read counts are estimated for each transcript using RSEM version 1.3.2 (22) and aggregated to obtain a quantification at the gene level.

### Single-cell dataset and gene expression

For the single-cell dataset composed of 10 PBMC cells type selected by FACS and sequenced separately (13). We downloaded the gene expression matrices for each cell types, from 10 datasets named as: CD8+ Cytotoxic T cells, CD8+/CD45RA+ Naïve Cytotoxic T Cells, CD56+ Natural Killer Cells, CD4+ Helper T Cells, CD4+ Helper T Cells, CD4+/CD45RO+ Memory T Cells, CD4+/CD45RA/CD25-Naïve T cells, CD4+/CD25+ Regulatory T Cells, CD34+ Cells, CD19+ B Cells, CD14+ Monocytes; available on 10X genomics web site (https://support.10xgenomics.com/single-cell-gene-expression/datasets), generate by Cell Ranger using the pipeline 1.1.0. Next, we merge these 10 matrices, to create the single-cell dataset of ∼95,000 cells.

To assess the reproducibility of each method in distinguishing each cell type from all others, we generated 20 replicates datasets that mimic typical single-cell experiments. Each replicate consisted of ∼10,000 cells, sampled from a normal distribution centered at 10,000 with a standard deviation of 1,000. Additionally, to assess the influence of cohort size on reproducibility, we repeated the experimental setup using datasets with varying mean cell counts—3,000, 5,000, and 8,000 cells.

### DEG and PG analyses

DEG results presented in this article were obtained using R version 4.0.0 and software packages MAST 1.14.0, limma 3.44.3, DESeq2 1.28.1, and edgeR 3.30.0. For each software, we used the standard pipeline described in their documentation, with default parameters. PG results were obtained using Python 3.11.5 and EPCY 0.2.6.4. All figures have been plotted using seaborn 0.10.1 and matplotlib 3.3.0 Python libraries. All analyses and figures in the paper are fully reproducible through scripts and data available in git repositories (https://github.com/iric-soft/epcy, https://github.com/iric-soft/epcy_paper) and zenodo (DOI:10.5281/zenodo.11639957, DOI:0.5281/zenodo.16507940, DOI:10.5281/zenodo.16508797).

To evaluate the robustness and specificity of DEG and PG methods, we conducted three complementary analyses across bulk and single-cell RNA-seq datasets. For bulk RNA-seq, subsampling experiments (Figure 2) were performed on the Leucegene3 dataset to assess the stability of statistical significance across varying sample sizes. DESeq2, edgeR, limma-voom, and EPCY were applied to multiple subsampling cohorts, and gene-level metrics were aggregated across replicates.

In Figure 3, we compared DEG and PG gene selection strategies using full bulk RNA-seq data. Genes were ranked by adjusted p-values or predictive scores, and selection thresholds were defined using quantiles of the full gene set. Overlaps and unique selections were visualized to highlight method-specific biases and biological relevance.

Figures 4 and 5 extended this comparison to single-cell RNA-seq data. Ten immune and hematopoietic cell types were analyzed across 20 replicates and four subsampling depths (3,000 to 10,000 cells). EPCY, Wilcoxon, MAST, and limma-trend were applied to each replicate, and reproducibility was assessed via volcano plots, pairwise replicate comparisons, and hexbin density plots. Quantile-based selection was used to evaluate consistency and specificity of gene detection across methods and conditions.

### Kernel Density Estimation and bandwidth

EPCY evaluates each gene’s predictive capacity through a cross-validated protocol. For each gene and each sample, we train a KDE classifier (C_kde_) for each condition. The KDE classifier uses a gaussian kernel and its bandwidth is estimated using the Scott’s variation of the Silverman’s rule of thumb (23), with a minimum threshold of 0.1 (by default).

Then C_kde_ is used to compute the probability of each class of a given sample.

Next, we draw *n* floats between 0 and 1 to assign *n* predicted class for each sample according to the probability computed with C_kde_, to fill *n* contingency tables (by default *n*=100). A Matthew’s Correlation Coefficient (MCC) is computed for each contingency table and is reported as a mean score and a 90% Confidence Interval (CI), by removing 5% of extreme values on both sides. Similarly to MCC, EPCY also computes mean PPV and NPV scores with their corresponding CI.

### Predictive scores

In this study, we use three predictive scores, computed with the number of samples being true positives (TP), true negatives (TN), false positives (FN), and true negatives (TN) reported in the contingency tables:

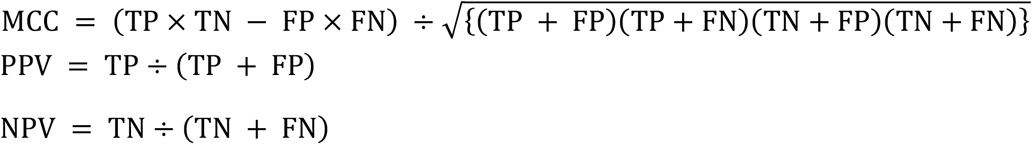

### EPCY’s reproducibility

To fill a contingency table, EPCY draws random values to assign a class according to probabilities learned by the KDE classifier (see Fig. 2). This means that different runs of EPCY can produce slightly different results. However, EPCY’s output is relatively stable, as each predictive score returned is a mean over several predictive scores, by default 100, which minimizes the variance between runs.

To ensure reproducibility, we added a parameter to fix the random seed. For all EPCY analyses shown in this article, the random seed was set to 42.

## ACKNOWLEDGEMENTS

The authors wish to thank Muriel Draoui for management of the Leucegene project and members of Institute for research in immunology and cancer (IRIC) genomic and bioinformatic platforms for support in data generation and analysis. The authors also acknowledge the contribution of the BCLQ (Banque de cellules leucémiques du Québec) staff (Giovanni D’Angelo, Claude Rondeau, Sylvie Lavallée) to this study. This work was supported by the Government of Canada through Genome Canada and the Ministère de l’économie et de l’innovation du Québec through Génome Québec (grant 4524 and grant 13528). Support from the Leukemia Lymphoma Society and the Canadian Cancer Society Research Institute to G.S. is also acknowledged. J.H. and G.S. hold a research chair from Industrielle-Alliance at Université de Montréal and a Bégin-Plouffe chair in blood stem cell chemogenomics of the Faculty of Medicine of Université de Montréal, respectively. BCLQ is supported by grants from the Cancer Research Network of the Fonds de recherche du Quebec–Santé. J.-F.S. is supported by an IVADO and Canada First Research Excellence Fund (Apogee/CFREF) postdoctoral award and a Canadian Institutes of Health Research fellowship (MFE-158159).

## AUTHORS CONTRIBUTIONS

Programming and algorithm development: É.A.

Methodology development: É.A, S.L.

Bulk RNA analyses: É.A, J-F S.

Single cell RNA analyses: É.A, V-P.L, J-F S.

Resources (primary human AML specimens): J.H.

Writing – Original Draft: É.A.

Writing – Review & Editing: S.L, J-F.S.

Funding acquisition: G.S, J.H, S.L.

