## Supplemental Figure S1 for "Predictive Gene Discovery with EPCY: A Density-Based Alternative to DE analysis"

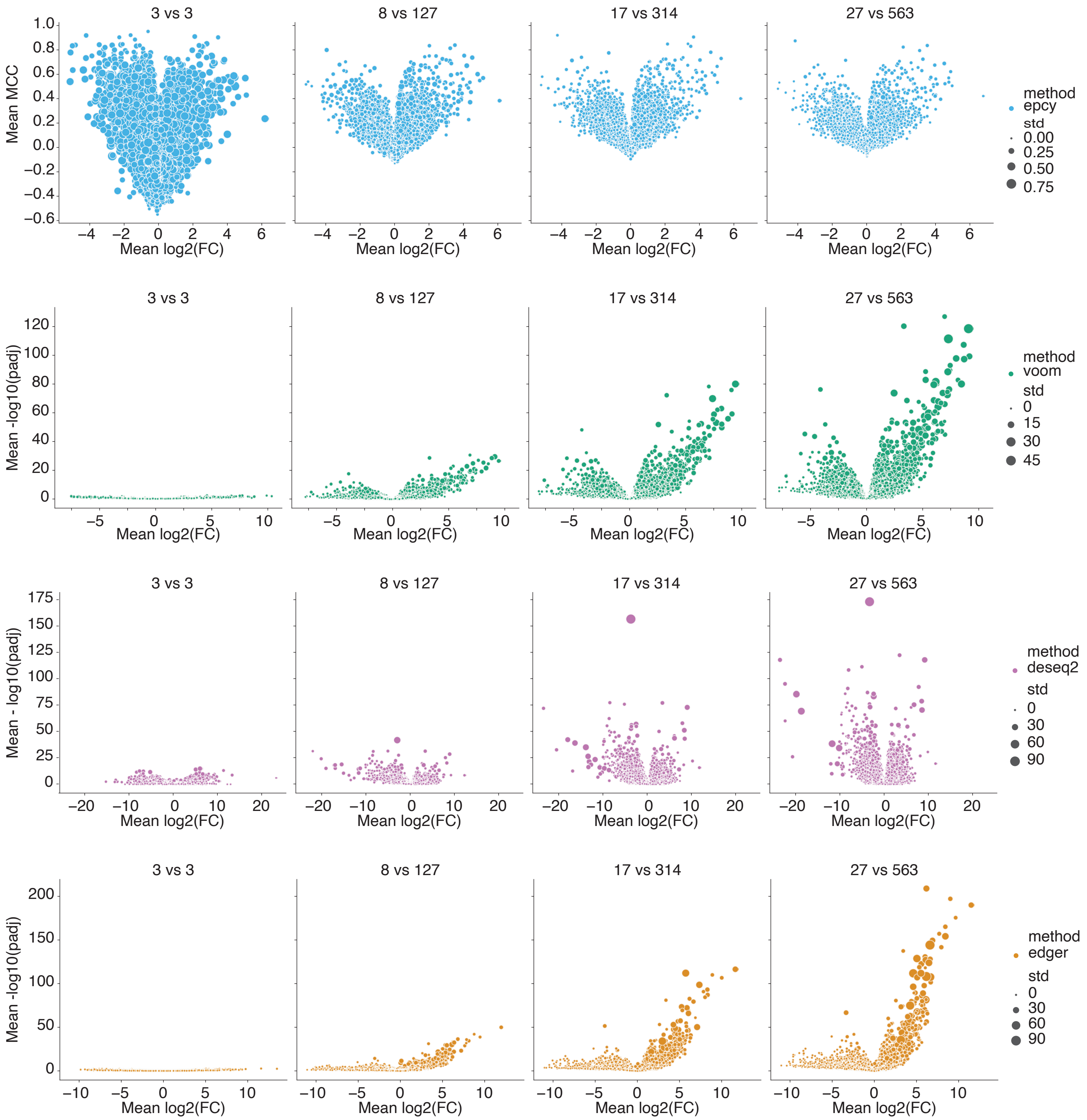


Figure S1: Volcano plots obtained by analysing t(15;17) subgroup of Leucegene cohort according to the number of samples used. For each size of the Leucegne cohort randomly sub-sampled, we create 10 replicates. The size of each plot represent the standard deviation of MCC or padj computed for each gene, using 10 replcates.
