## Supplemental Figure S2 for "Predictive Gene Discovery with EPCY: A Density-Based Alternative to DE analysis"

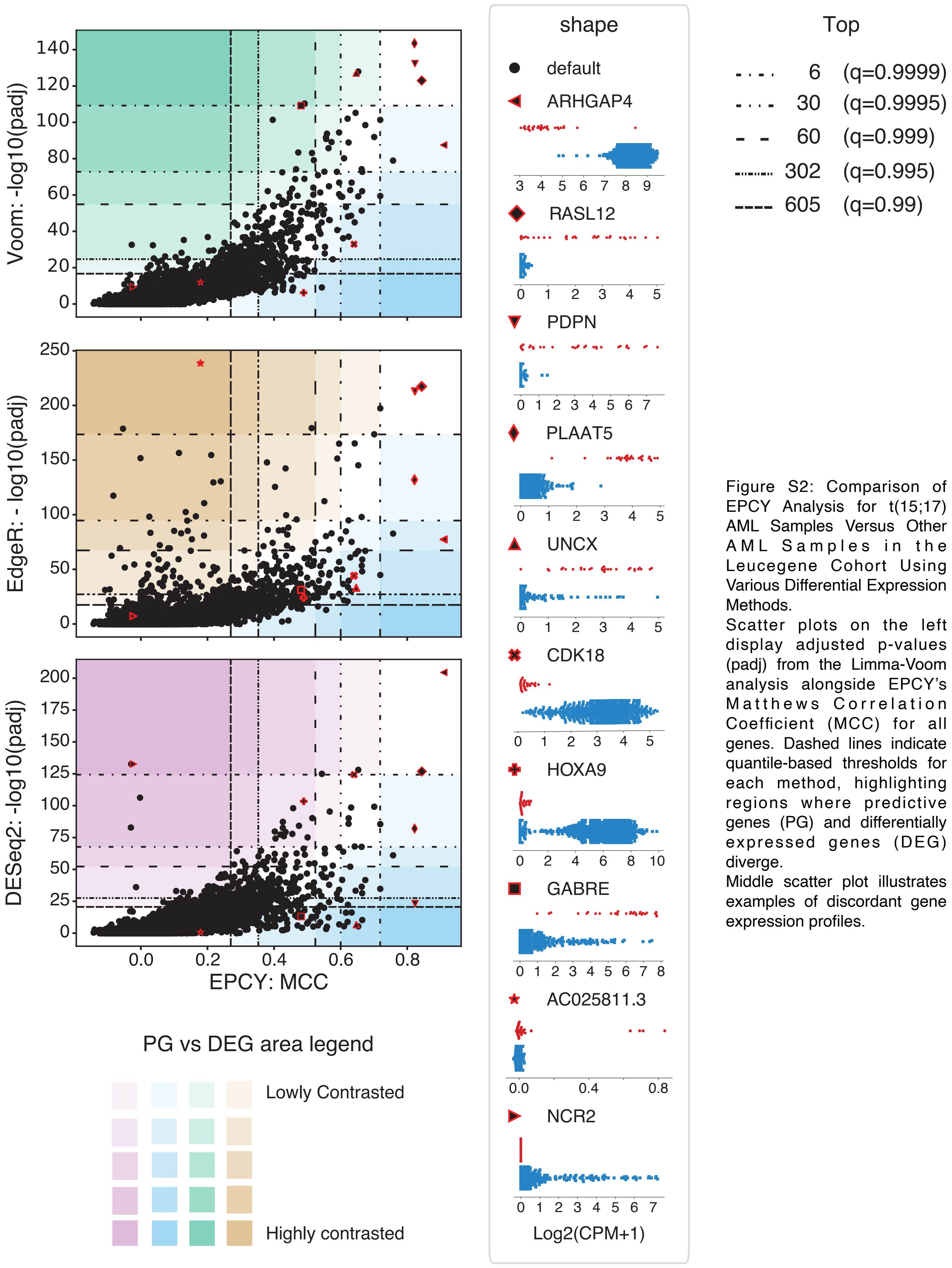


Figure S2: Comparison of EPCY Analysis for t(15;17) AML Samples Versus Other AML Samples in the Leucegene Cohort Using Various Differential Expression Methods.
Scatter plots on the left display adjusted p-values (padj) from the Limma-Voom analysis alongside EPCY's Matthews Correlation Coefficient (MCC) for all genes. Dashed lines indicate quantile-based thresholds for each method, highlighting regions where predictive genes (PG) and differentially expressed genes (DEG) diverge.
Middle scatter plot illustrates examples of discordant gene expression profiles.
