## Supplemental Figure S3 for "Predictive Gene Discovery with EPCY: A Density-Based Alternative to DE analysis"

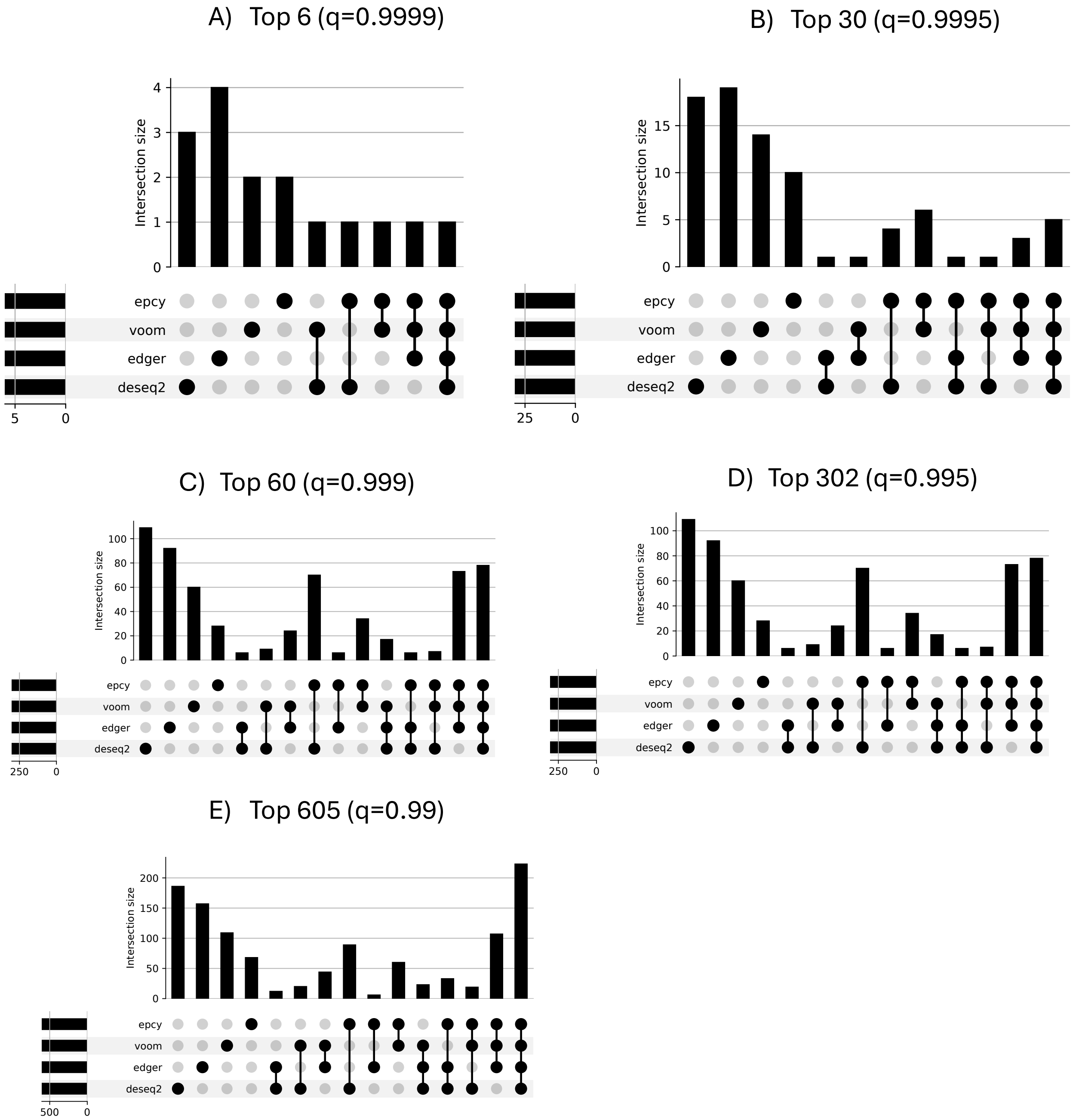


Figure S3: The number of intersecting genes for each method according to thresholds, using analyses of t(15;17) AML samples of the Leucegene cohort. Empty intersections are omitted.
