## Supplemental Figure S4 for "Predictive Gene Discovery with EPCY: A Density-Based Alternative to DE analysis"

A) Top 6 (q=0.9999)

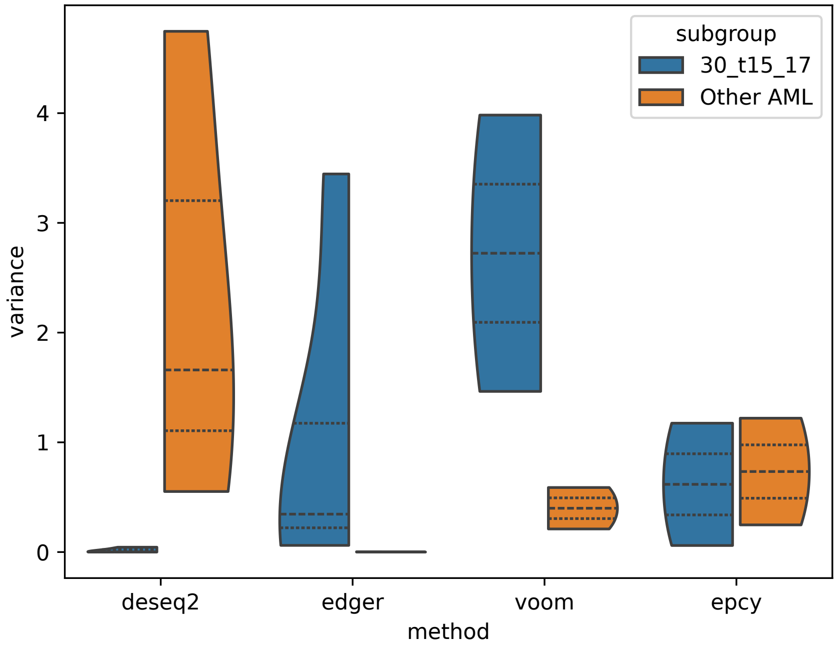


B) Top 30 (q=0.9995)

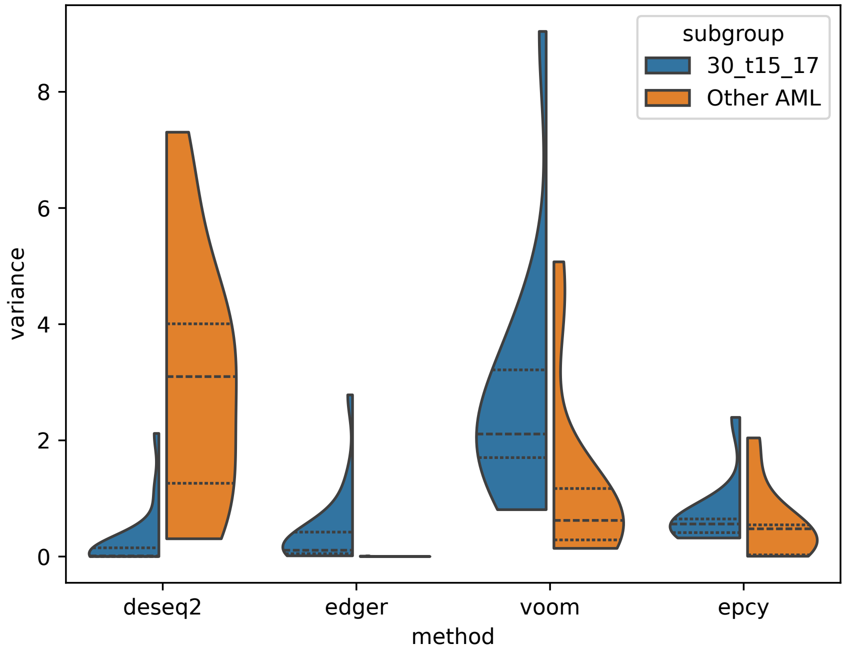

C)  Top 60 (q=0.999)

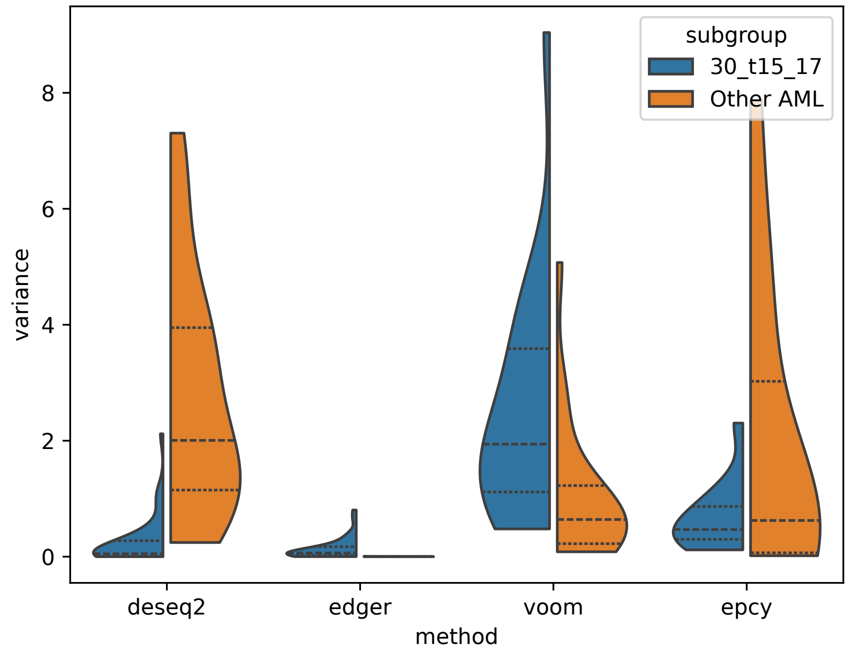


D)  Top 302 (q=0.995)

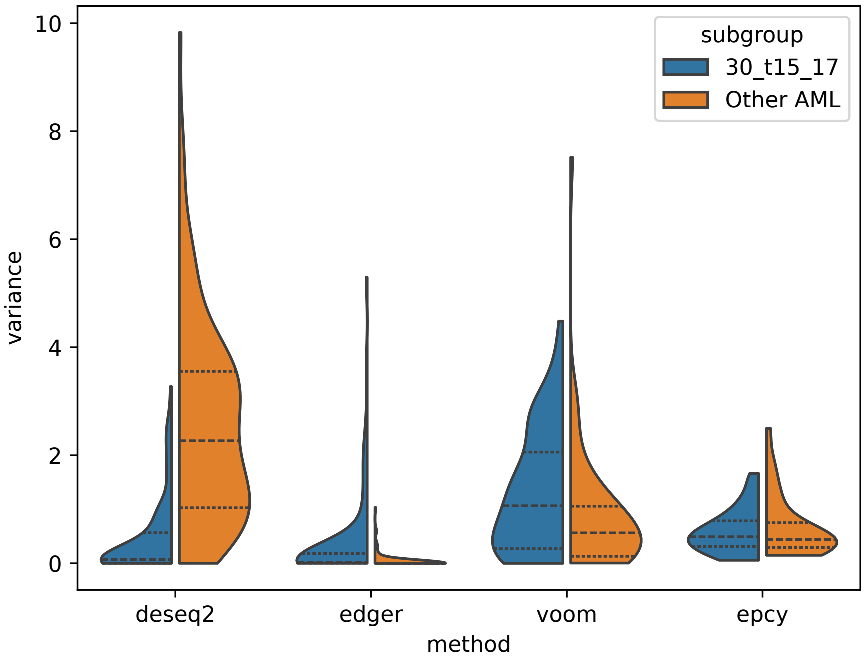

E)   Top 605 (q=0.99)

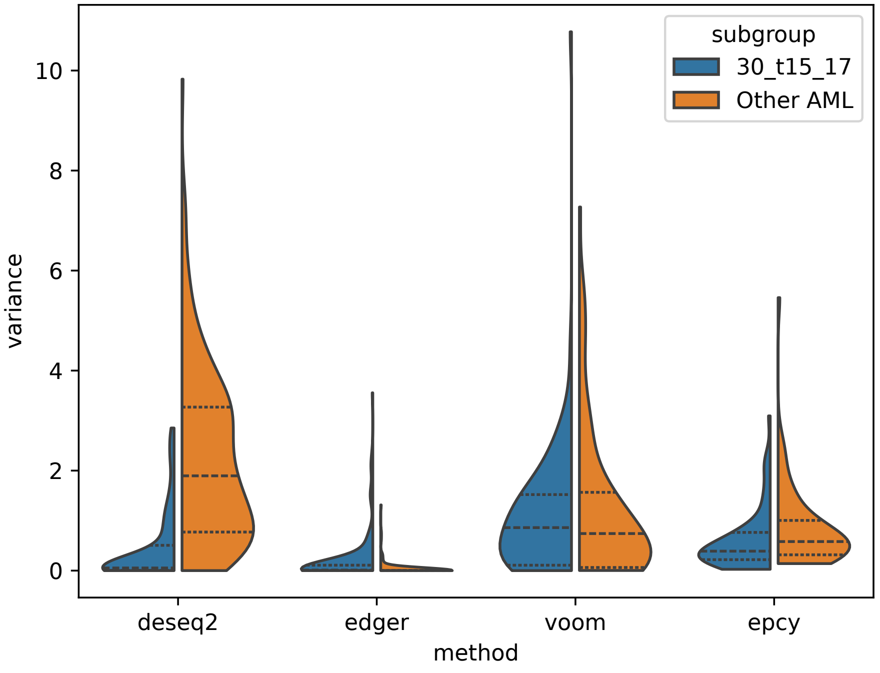


Figure S4: Using t(15;17) AML samples of the Leucegene cohort, this figure presents the density of variance for mutually exclusive selections of genes, according to thresholds selected. Quartiles are indicated by vertical dashed lines.
