## Supplemental Figure S5 for "Predictive Gene Discovery with EPCY: A Density-Based Alternative to DE analysis"

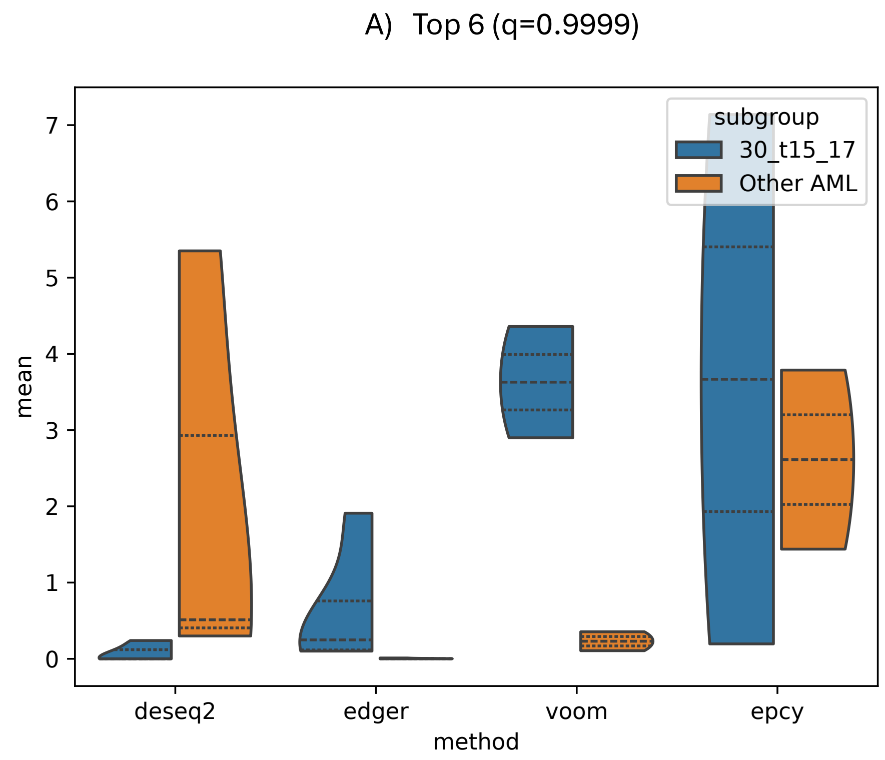
A) Top 6 (q=0.9999)
subgroup
30_t15_17
Other AML
mean
0
1
2
3
4
5
6
7
deseq2
edger
voom
epcy
method

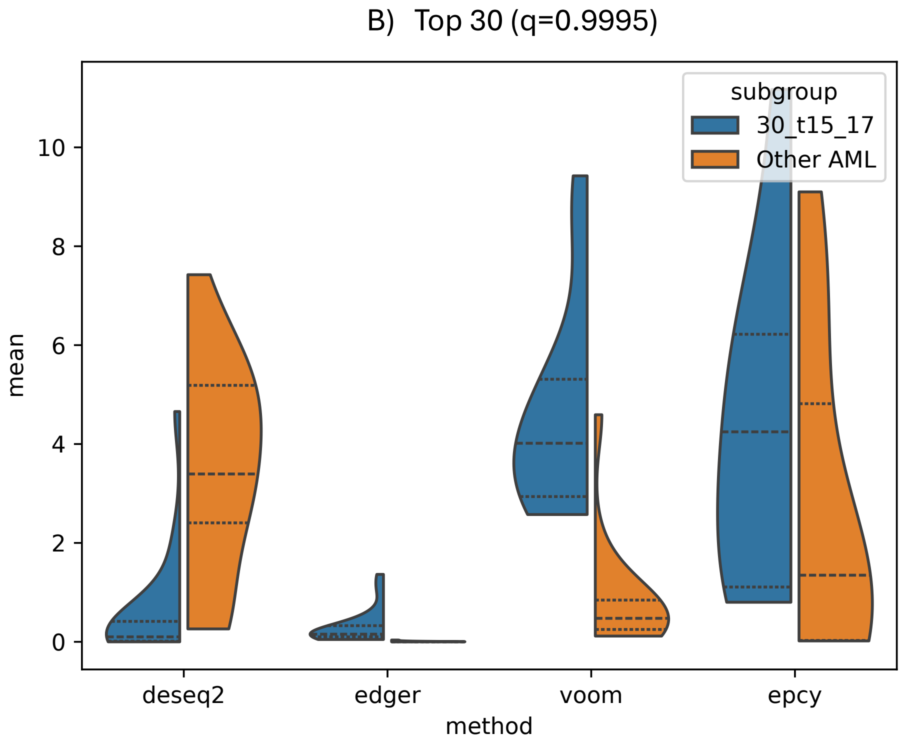
B) Top 30 (q=0.9995)
subgroup
30_t15_17
Other AML
mean
0
2
4
6
8
10
deseq2
edger
voom
epcy
method

C) Top 60 (q=0.999)

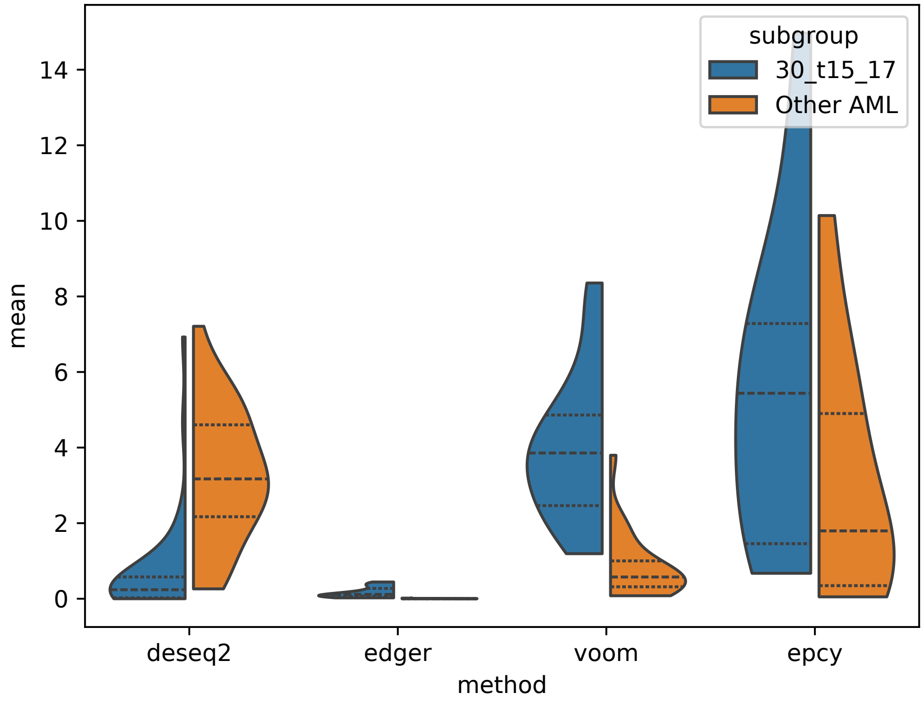


D) Top 302 (q=0.995)

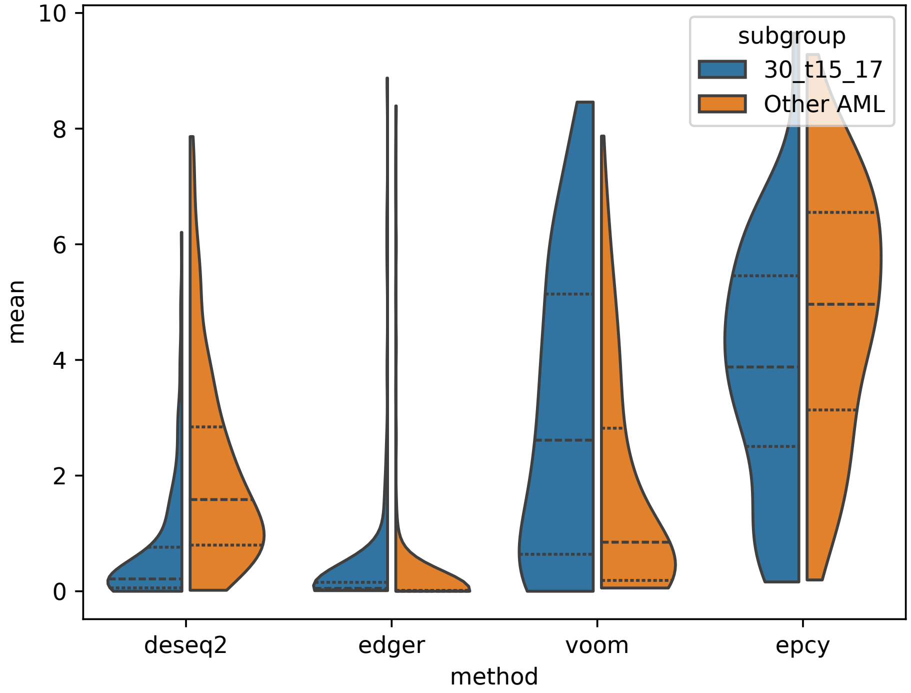

E) Top 605 (q=0.99)

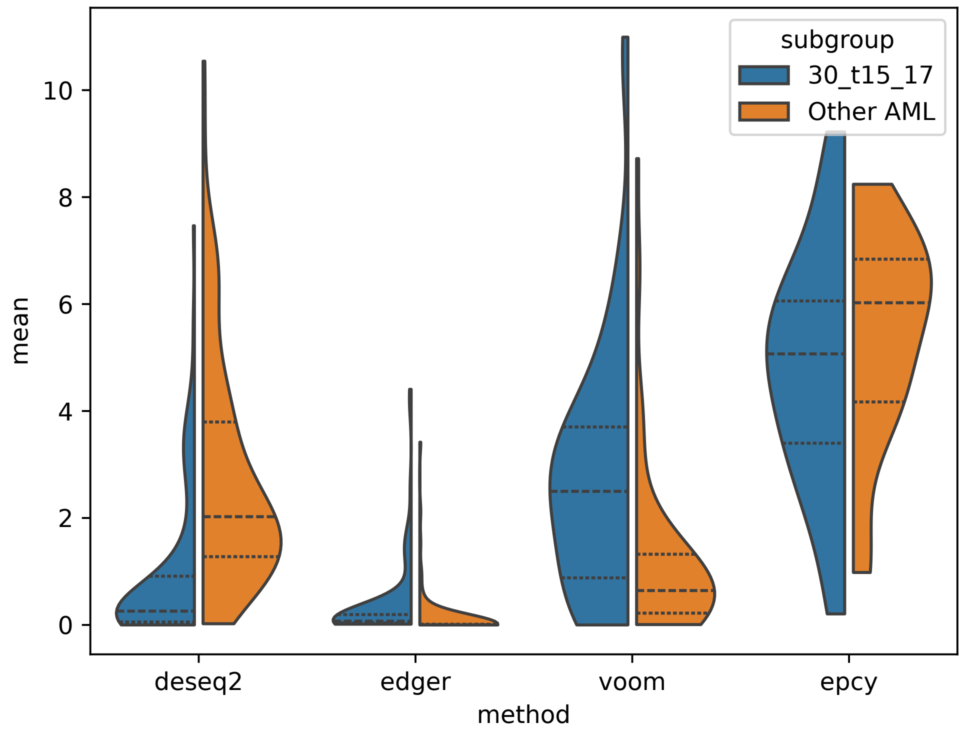


Figure S5: Using t(15;17) AML samples of the Leucegene cohort, this figure presents the density of mean expression for mutually exclusive selections of genes, according to thresholds selected. Quartiles are indicated by vertical dashed lines.
