## Supplemental Figure S6 for "Predictive Gene Discovery with EPCY: A Density-Based Alternative to DE analysis"

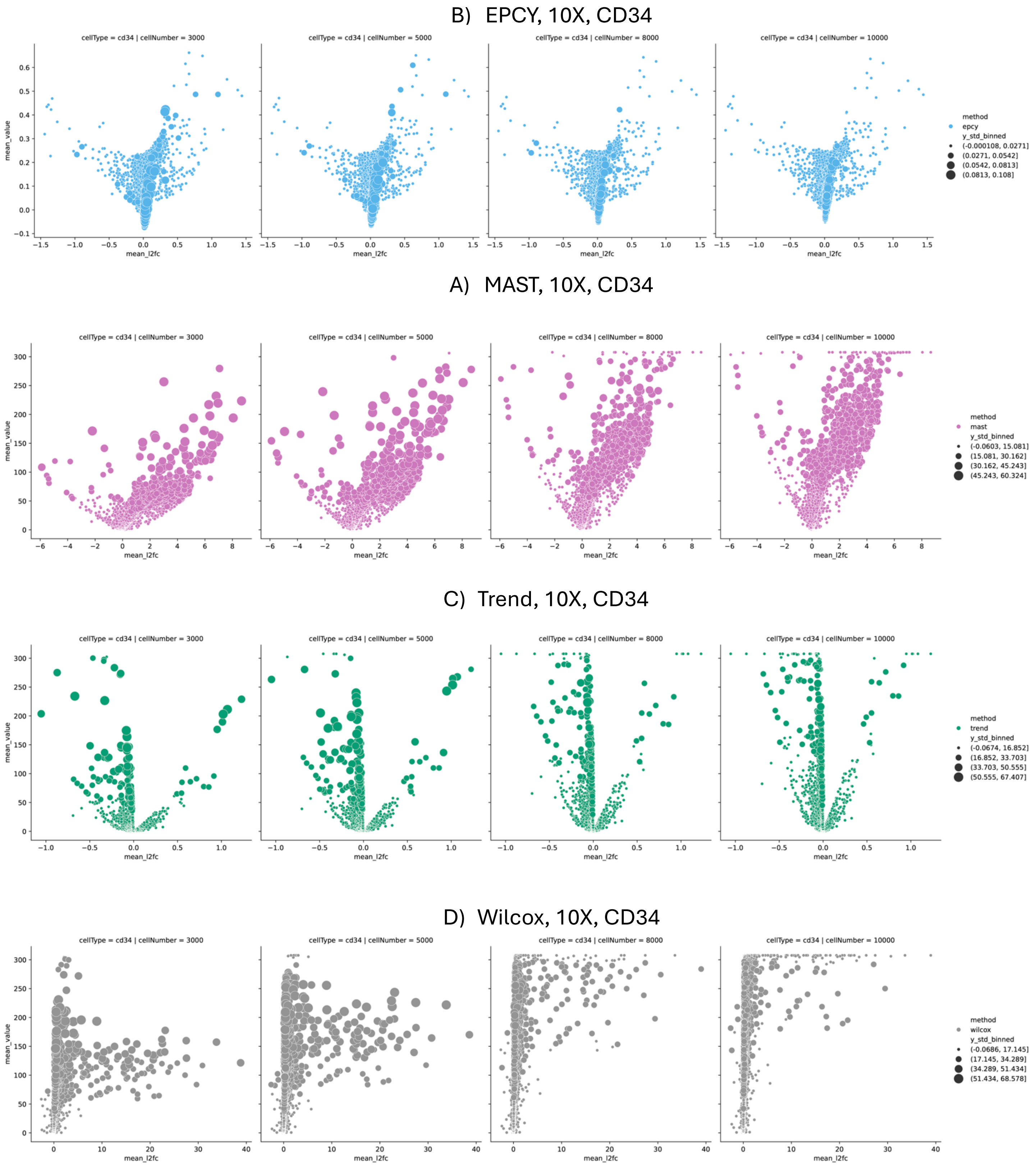


Figure S6: Volcano plots illustrate the relationship between MAST, Trend and Wilcox adjusted p-values (padj) and their standard deviations across the 20 replicates, highlighting how both metrics increase with cohort size. In contrast, EPCY's Matthews Correlation Coefficient (MCC) remains largely stable, showing minimal variation regardless of the number of cells.
