## Supplemental Figure S7 for "Predictive Gene Discovery with EPCY: A Density-Based Alternative to DE analysis"

11 vs 18

3,000 cells

10,000 cells

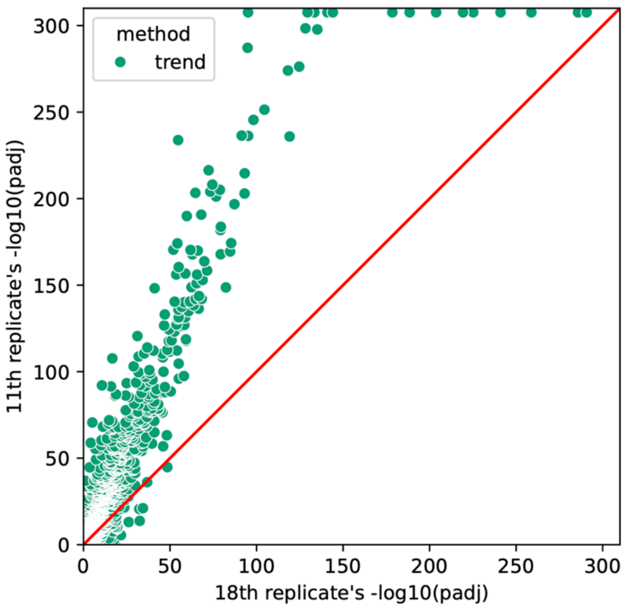


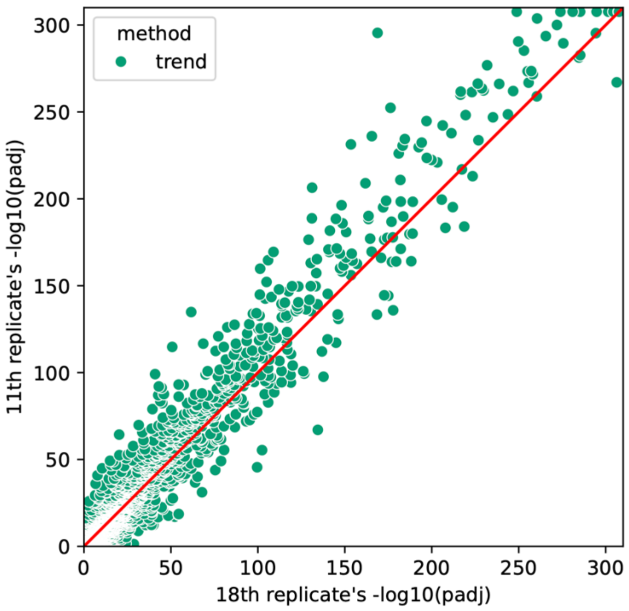


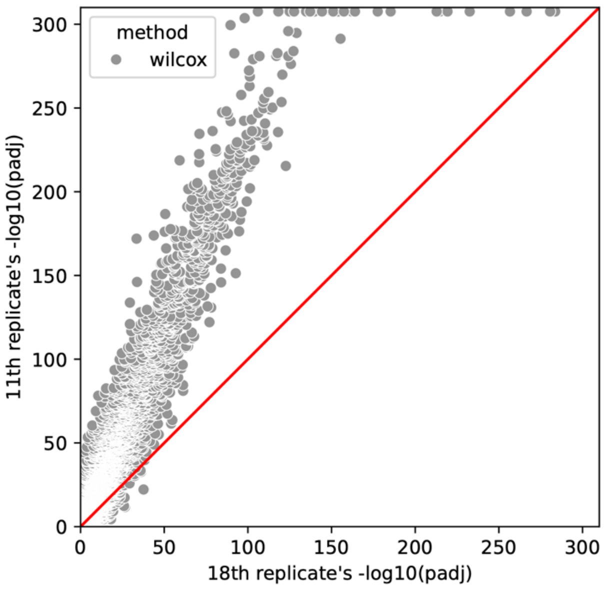


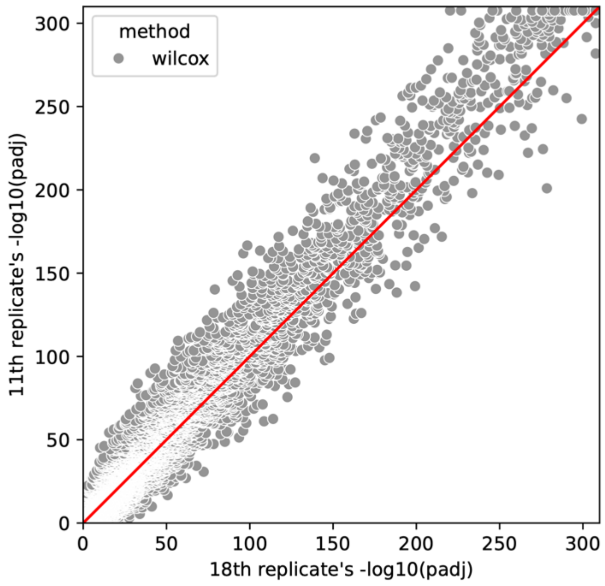

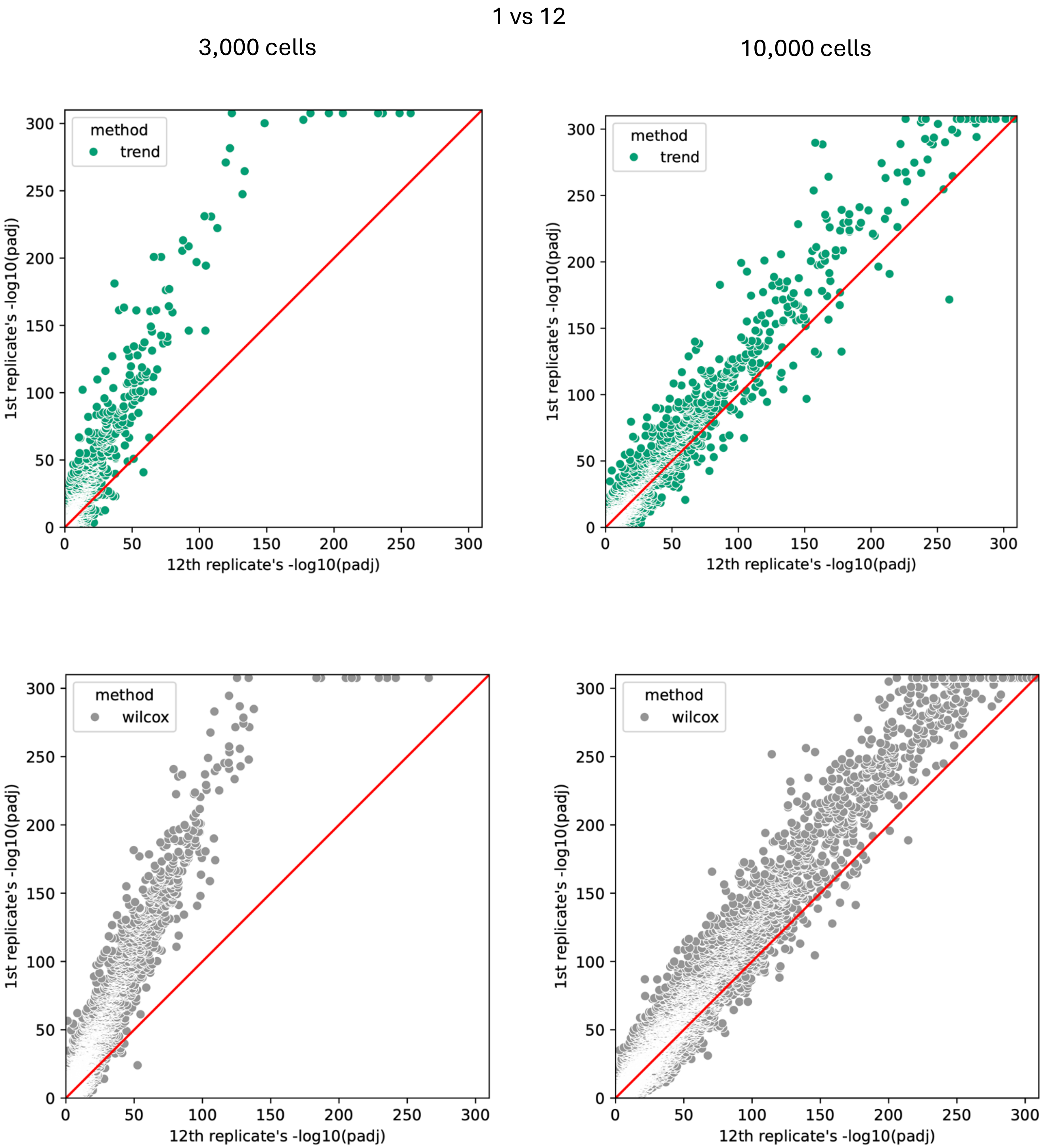


Figure S7: Scatter plots of Trend and Wilcox adjusted p-values (padj) resulting from the analysis of two pairs of replicates constituted of 3,000 and 10,000 cells, respectively. The red diagonal line indicates the expected trend for perfect reproducibility between datasets.
