## Supplemental Table S1 for "Predictive Gene Discovery with EPCY: A Density-Based Alternative to DE analysis"

Supplementary Table 1: Computer ressources need for each methods, in fonction of number of samples analysed

| | 3 vs 3 | 6 vs 65 | 8 vs 127 | 11 vs 190 | 14 vs 252 | 17 vs 314 | 19 vs 376 | 22 vs 501 | 25 vs 501 | 27 vs 563 |
|---|---|---|---|---|---|---|---|---|---|---|
| **EPCY (4 threads)** | 00:01:31 | 00:10:47 | 00:17:18 | 00:29:44 | 00:42:44 | 00:47:24 | 00:50:51 | 00:58:55 | 01:06:21 | 01:14:22 |
| | 2.6Gb | 2.7Gb | 2.8Gb | 2.8Gb | 3.4Gb | 3.7Gb | 3.9Gb | 4.1Gb | 4.3Gb | 4.4Gb |
| **Limma Voom** | 00:00:33 | 00:00:46 | 00:00:55 | 00:01:44 | 00:01:09 | 00:00:54 | 00:01:00 | 00:01:02 | 00:01:16 | 00:01:24 |
| | 0.7Gb | 0.7Gb | 0.9Gb | 1.5Gb | 1.2Gb | 1.3Gb | 1.4Gb | 2.3Gb | 2.5Gb | 3.0Gb |
| **DESeq2 (4 threads)** | 00:01:02 | 00:01:50 | 00:01:41 | 00:01:57 | 00:02:35 | 00:02:46 | 00:03:23 | 00:03:36 | 00:04:37 | 00:04:31 |
| | 1.2Gb | 6.2Gb | 6.6Gb | 6.7Gb | 8.7Gb | 9.6Gb | 10.8Gb | 12.3Gb | 14.4Gb | 13.4Gb |
| **EdgeR** | 00:00:33 | 00:01:28 | 00:01:40 | 00:02:21 | 00:04:03 | 00:03:44 | 00:04:26 | 00:05:08 | 00:07:32 | 00:08:11 |
| | 0.7Gb | 0.9Gb | 1.0Gb | 1.3Gb | 1.6Gb | 2.1Gb | 2.3Gb | 2.5Gb | 2.8Gb | 3.17Gb |
